# Cingulate dependent social risk assessment in rats

**DOI:** 10.1101/452169

**Authors:** Yingying Han, Rune Bruls, Rajat Mani Thomas, Vasiliki Pentaraki, Naomi Jelinek, Mirjam Heinemans, Iege Bassez, Sam Verschooren, Illanah Pruis, Thijs Van Lierde, Nathaly Carrillo, Valeria Gazzola, Maria Carrillo, Christian Keysers

## Abstract

Social transmission of distress has been conceived of as a one-way phenomenon in which an observer catches the emotions of another. Here we use a paradigm in which an observer rat witnesses another receive electroshocks. Bayesian model comparison and Granger causality argue against this one-way vision in favor of bidirectional information transfer: how the observer reacts to the demonstrator’s distress influences the behavior of the demonstrator. Intriguingly, this was true to a similar extent across highly familiar and entirely unfamiliar rats. Injecting muscimol in the anterior cingulate of observers reduced freezing in the observers and in the demonstrators receiving the shocks. That rats share the distress of unfamiliar strains is at odds with evolutionary thinking that empathy should be biased towards close individuals. Using simulations, we support the complementary notion that distress transmission could be selected to more efficiently detect dangers in a group.

## 1. Introduction

Empathy, the ability to share and understand the emotional state of other individuals is thought to be crucial for successful social interactions. Accumulating evidence suggests that rodents have at least a precursor of empathy called emotional contagion (for review see Sehoon Keum & Shin, 2016; K. Z. Meyza, Bartal, Monfils, Panksepp, & Knapska, 2017; Panksepp & Lahvis, 2011; Sivaselvachandran, Acland, Abdallah, & Martin, 2016). Whereas empathy proper requires the ability to distinguish emotions of the self from those vicariously shared on behalf of others, emotional contagion only requires that an observer’s emotional state gets to resemble that of a target without specific cognitive attributions (Bernhardt & Singer, 2012; de Waal & Preston, 2017; Preston & de Waal, 2002). Empathy, to be adaptive, is thought to be biased towards individuals that are socially and genetically closer to the empathizer: “Empathy is still subject to appraisals, filters and inhibitions that prevent it from being expressed when it would be maladaptive. […] the empathic response is increased by similarity, familiarity and social closeness … and this is consistent with where evolutionary theory expects it to occur: that is, in close interdependent social relationships that involve either genetic relatedness or reciprocation” (de Waal & Preston, 2017).

While empathy proper is perhaps difficult to study in rodents, emotional contagion in these animals has been the subject of a rapidly growing number of studies. Prototypical example of how emotional contagion is measured are experiments in which one animal receives a footshock, and the freezing of another that witnesses the event is found to be increased, suggesting that the distress of the shocked demonstrator was transferred to the observer (Atsak et al., 2011; Carrillo et al., 2015; Gonzalez-Liencres, Juckel, Tas, Friebe, & Brüne, 2014; Jeon et al., 2010; S. Keum et al., 2016; S. Kim, Mátyás, Lee, Acsády, & Shin, 2012; Sanders, Mayford, & Jeste, 2013). These experiments often investigate which factors influence the extent to which the witness ‘catches’ the emotion of the demonstrator. Due to its importance for regulating empathy, the effect of familiarity on emotional contagion has been extensively studied in mice (Gonzalez-Liencres et al., 2014; Jeon et al., 2010; Langford, Crager, Shehzad, & Smith, 2006; Martin et al., 2015), which shows that increasing the level of familiarity across demonstrators and observers increases how much the demonstrator influences the observer. This suggests that even if spontaneous, emotional contagion is also regulated by factors regulating empathy, such as familiarity, which led many to consider emotional contagion as a pre-cursor of empathy. Is the same true for rats? In contrast to mice no studies have tested the role of familiarity in the behavioral response of rats *directly* witnessing a conspecific experience a painful stimulus. What we do know is that interactions with a conspecific that had been exposed to a painful stimulus *elsewhere* can lead to stronger effects in more familiar individuals (Li et al., 2014) or in *siblings* (Jones, Riha, Gore, & Monfils, 2014). However, since the imminence of a threat changes the behavioral and neural responses of an animal(Fanselow, 1994), transmission of a state influenced by past danger signals (potentially via olfactory cues) differs from witnessing an acute reaction to distress (partially via auditory and visual cues). Finally, rats will help trapped individuals from a strain they are familiar with more than animals from a strain they are unfamiliar with (Ben-Ami Bartal, Rodgers, Bernardez Sarria, Decety, & Mason, 2014), but such prosocial behavior may be more tightly regulated due to its potential cost than emotional contagion.

Although social interactions are by nature bidirectional, emotional contagion is usually studied, both in animals and humans, as the transfer of emotion in a unidirectional manner from one individual in which an experimental manipulation triggers an emotion (the demonstrator) to another that is made to witness the event (the observer, Figure 1A). Does it make sense to conceive of emotional contagion as a one-way information transfer? Indirect evidence against this notion comes from a related but largely distinct field investigating social buffering: the emotional reaction of a stressed animal is sometimes attenuated when surrounded by non-stressed bystanders (Davitz & Mason, 1955; Fuzzo et al., 2015; Guzmán et al., 2009; Ishii, Kiyokawa, Takeuchi, & Mori, 2016; Kikusui, Winslow, & Mori, 2006; Kiyokawa, Hiroshima, Takeuchi, & Mori, 2014; Kiyokawa, Honda, Takeuchi, & Mori, 2014; Mikami, Kiyokawa, Takeuchi, & Mori, 2016; Terranova, Cirulli, & Laviola, 1999) (Figure 1B). More generally, a growing number of scientists argue that if we wish to understand the nature of social *interactions* we must develop methods and paradigms that can study *bidirectional* influences across individuals in face-to-face situations rather than simply exposing subjects to prerecorded stimuli (Schilbach et al., 2013). A successful example of how focusing on inter-individual interactions can generate conceptual advances comes from cowbirds. Only once real-time interactions across males and females were studied and experimentally manipulated did we get to understand that males learn to perform attractive songs using interactive feedback from the female: the males sing to females, the females then signal back how much they like that particular song by flapping their wings, and the male uses this feedback to shape the song towards the most attractive variants (White, 2010).

**Figure 1.**
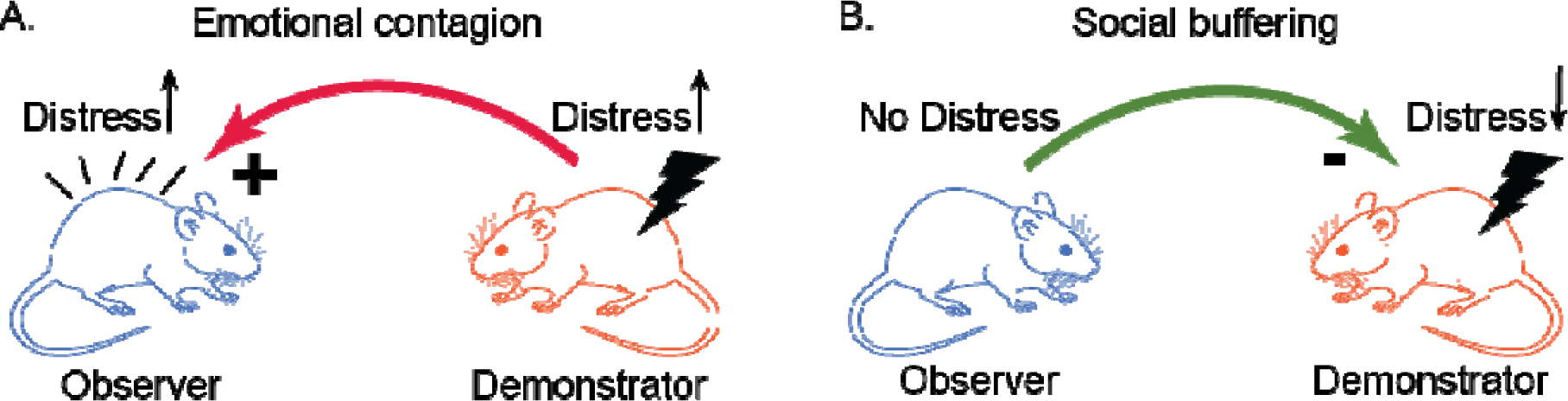
Emotional contagion and Social buffering paradigms. (A) A schematic representation of the paradigm used to investigate emotional contagion. An observer rat witnesses a demonstrator rat receive an electric foot shock. The shock induces fear and pain responses in the demonstrator, which in turn is unidirectionally transferred to the observer. In these paradigms, the variable of interest is the amount of distress of the observer. B) A schematic representation of the social buffering paradigm. A demonstrator rat receives an electric footshock. The fear response of the demonstrator is reduced in the presence of an observer rat. The variable of interest is the amount of distress of the demonstrator.

To shed further light onto the nature of emotional contagion as a channel of social *interactions*, we will therefore take stock of these observations and address two questions. First whether a *mutual* influence across individuals offers a better explanation of the behavior of rats in a prototypical emotional contagion paradigm. Second, whether familiarity has the strong modulatory effect on this phenomenon in rats that would be expected for empathy. For both questions, we harness a paradigm we developed in the lab in which a shock-experienced observer rat interacts through a perforated transparent divider with a demonstrator rat receiving footshocks. We quantify the freezing behavior of both animals during an initial 12 minute baseline period and a 12 minute test period in which the demonstrator received 5 footshocks (1.5mA, 1s each, ISI: 240-360s, Figure 2).

**Figure 2.**
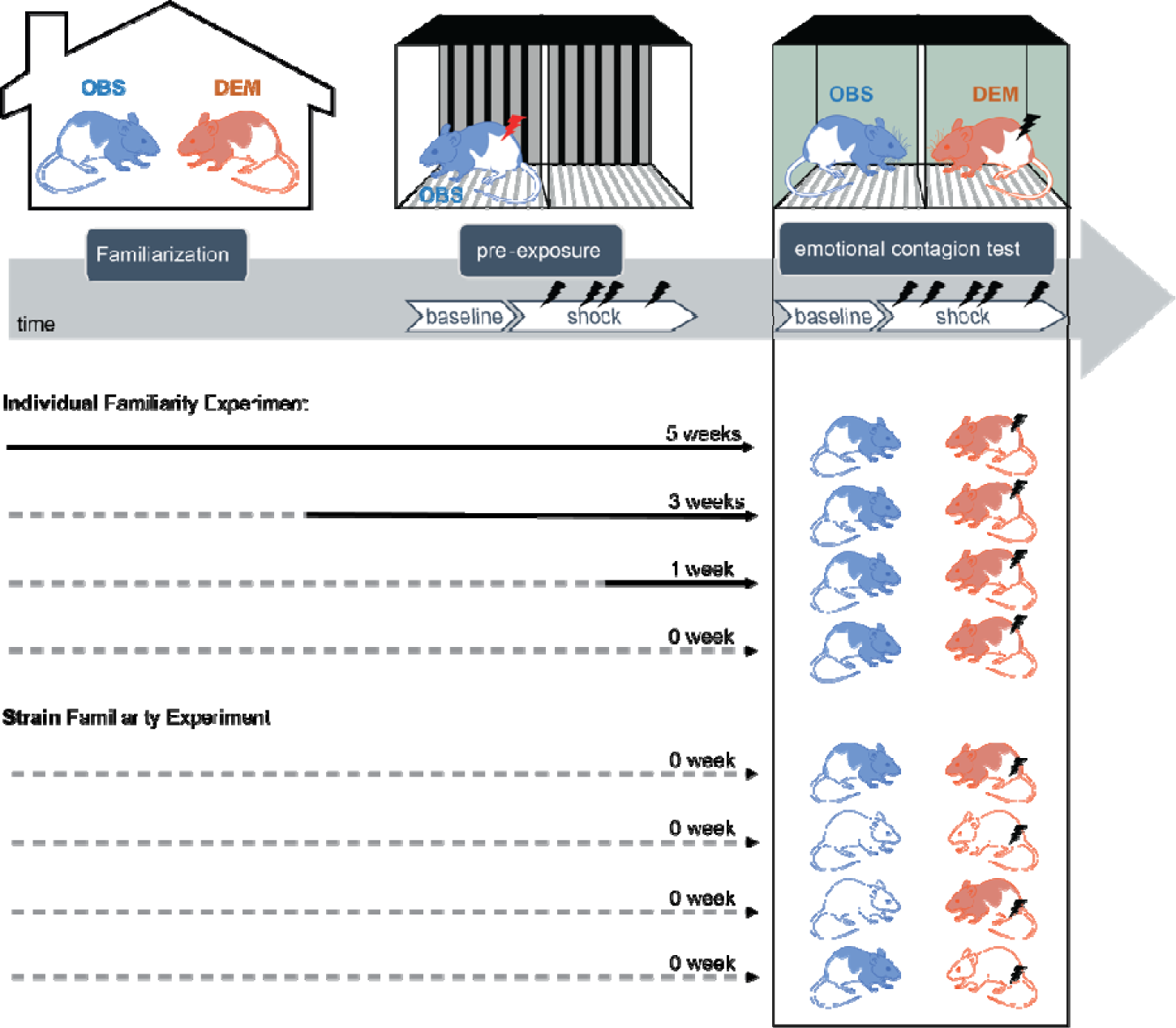
Experimental procedure. The procedure started with a familiarization phase (top left) in which a demonstrator rat (DEM in orange) was housed together with an observer rat (OBS in blue) for different periods of time depending on the experimental groups. The time the dyad spent together before test day (middle and bottom panel) is depicted as a solid time line (for 1, 3 or 5 weeks) while the dashed sections of the lines indicate periods during which the demonstrator and observer were housed apart. After the familiarization phase, the OBS were pre-exposed to footshocks alone (top middle). The pre-exposure procedure consisted of a 12 minute baseline and a 12 minute shock period in which the observer received 4 footshocks (0.8mA, 1s each, ISI: 240-360s). This was followed by the emotional contagion test (top right) consisting of a 12 minute baseline and a 12 minute shock period. During the shock period the observer witnessed the demonstrator in an adjacent chamber receive 5 footshocks (1.5mA, 1s each, ISI: 120-180s). In experiment 1 all animals were Long Evans (hooded rats in the middle panel). In experiment 2, the demonstrator-observer dyads were either from the same strain (i.e. both hooded Long Evans or both albino Sprague Dawley) or from different strains (i.e. one hooded Long Evans and one albino Sprague Dawley).

To address our first question, we leverage the fact that in our paradigm the demonstrator can witness the observer’s reaction, and use Bayesian model fitting, model comparison and Granger causality to investigate whether the freezing of the demonstrator also reflects feedback influences from the observer’s reaction. We predict there is feedback flow of information. To address our second question, we manipulated familiarity in two ways. In the first experiment (Individual Familiarity Exp), all demonstrator-observer dyads were from the same strain (i.e. Long Evans) but differed in how long they had been housed together with that particular individual. In the second experiment (Strain Familiarity Exp), all demonstrator-observer dyads were unfamiliar with the animal they were paired with during the emotional contagion experiment but differed in whether rats were familiar with the strain of their pair-mate (i.e. both Long Evans or both Sprague Dawley) or were unfamiliar with that strain (i.e. one Long Evans and one Sprague Dawley). Our prediction, based on mice studies, is that less familiar animals would show reduced evidence of influence in both directions.

Finally, two follow-up experiments using pharmacological deactivations of the cingulate and behavioral simulations investigate the neural locus and value of the social transmission of distress. Our prediction is that deactivating the cingulate of observer animals would reduce their response and that this would feed-back to reduce the demonstrators’ response.

## 2. Results

### 2.1 Emotional contagion: general behavioral responses

The scatter plots of Figure 3 show, for both experiments, how much observers and demonstrators froze during the 12 minute baseline (when no shock was delivered, black dots) and during the 12 minute shock period during which the demonstrator received 5 shocks (red crosses). The elongated shape of the scatter plots during the shock period suggests a relationship between the freezing levels of demonstrators and observers: Dyads in which demonstrators froze most are often dyads in which the observers also froze most. To explore the directionality of this relationship we will use Bayesian modeling and Granger causality.

**Figure 3.**
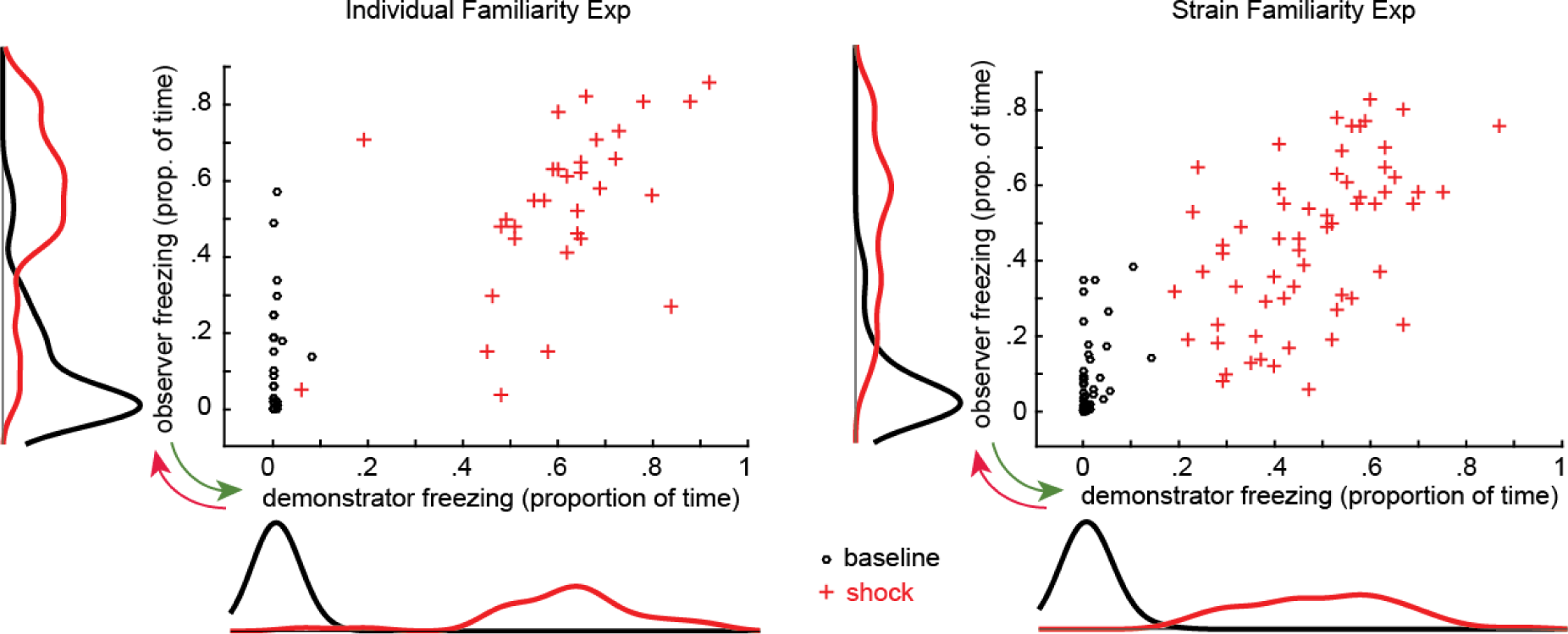
Observer and Demonstrator freezing responses during the emotional contagion paradigm. For both the Individual Familiarity and the Strain familiarity experiments the scatter plots indicate the proportion of time spent freezing by the demonstrator and observer both during baseline (black dots) and the shock period (red crosses). The marginal histograms indicate the distribution of freezing behavior during baseline (black lines) and the shock period (red lines). Red arrow: possible influence of the demonstrator freezing on the observer freezing (akin to as emotional contagion). Green arrow: possible influence of the observer freezing on the demonstrator freezing (akin to as social buffering if the level of the observer freezing is lower than that of the demonstrator).

### 2.2 Effect of Familiarity and Feedback – Bayesian Model Comparison

Bayesian modeling was used to (a) compare models with feedback, in which the freezing of the observer influences back how much the demonstrator freezes, against models without feedback; and (b) to identify whether the time individuals spent together influences the coupling between the animals. Separate models were constructed using different combinations of experimental variables that could explain the observer and demonstrator freezing in the two experiments. Figure 4 summarizes the variables included in the models, with those that significantly explain the observer’s and demonstrator’s freezing marked in red.

**Figure 4.**
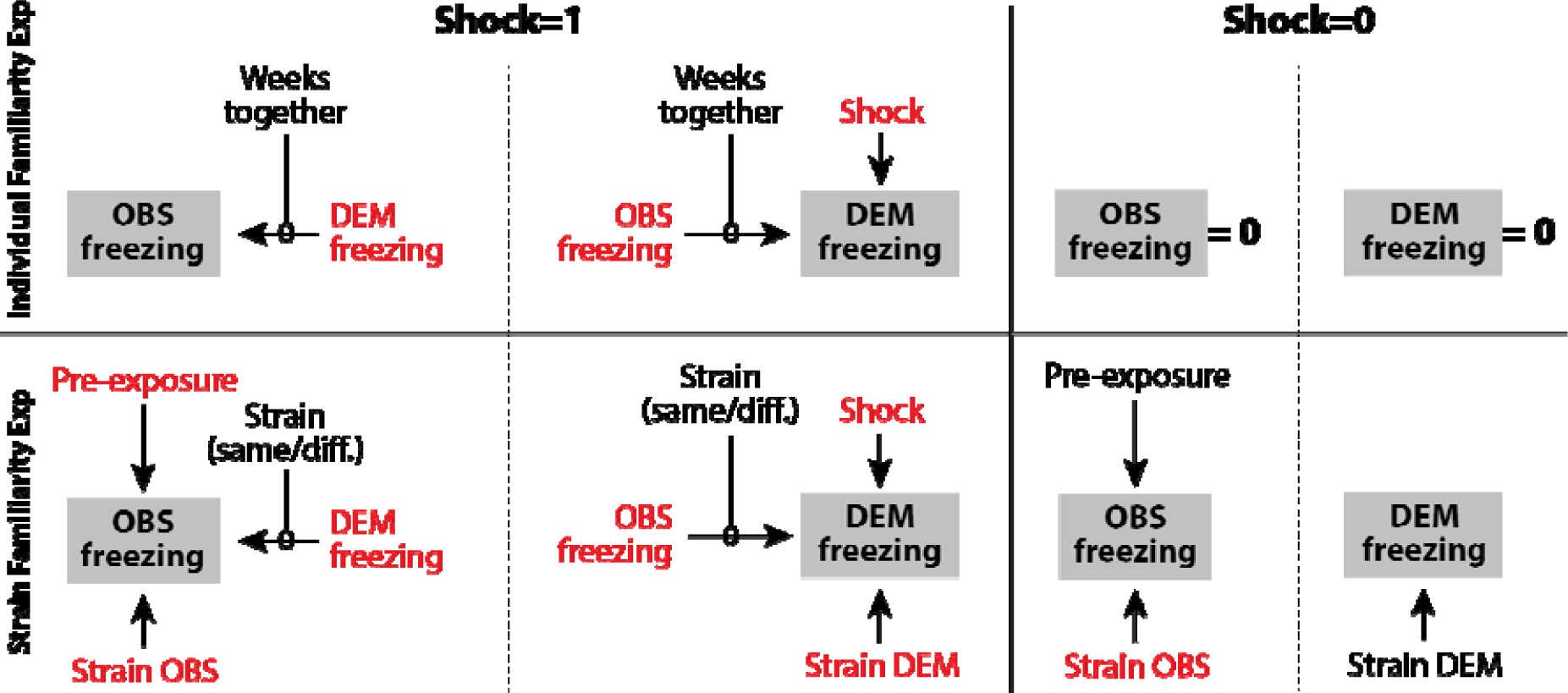
Variables included in the Bayesian models. Several models were built, separately for the two experiments, based on the factors that could describe the observer and demonstrator freezing. Here the full models for the Individual Familiarity Experiment (top) and Strain Familiarity Experiment (bottom) are shown separately for epochs in which shocks are delivered (Shock=1) and those in which no shocks are delivered (Shock=0). The target variable that the models explains are shown in a gray box. These full models were then compared against simplified models, and the variables included in the winning model are shown in red. The modulator ‘Weeks together’ captures whether the effect across animals depends on the number of weeks the observer and demonstrator spent together before testing (i.e. 0, 1, 3 and 5 weeks). This modulator was implemented in two different ways (see Table 2 and 3, and Figure 5): (i) in a way that models a linear increase of interindividual influence with number of weeks spent together, with the impact thus 5 times stronger after 5 compared to 1 week spent together or (ii) in a way that simply models a different connection weight for each group. Strain OBS and strain DEM capture the effect of a particular strain on the average freezing level of that strain. Strain (same/diff.) is a binary variable indicating whether the observer and demonstrator dyad were of the same or different strain. Finally, the variable pre-exposure indicates the amount of freezing of the observer during pre-exposure. Unfortunately, we only collected movies during pre-exposure in the Strain Familiarity Experiment, and thus cannot retrospectively include that variable in the models of the Individual Familiarity Experiment.

#### 2.2.1 Results from the Individual Familiarity Experiment

##### Demonstrator’s freezing

Of the eight tested models, the one best explaining the demonstrator’s freezing shows that Freezing_dem_ = 0.39 x Shock_dem_ + 0.41 x Freezing_obs_ x Shock_dem_ (*model 6*, elpd_loo_ estimate = 45.3 and SE = 16.1; Table 1A). This indicates that within our paradigm the freezing of the demonstrator (Freezing_dem_) is approximated by assuming that it is zero when no shock is delivered (since the variable Shock_dem_ is then equal to zero, nulling all elements of the equation). However, when a shock is delivered, the demonstrator’s freezing can be estimated at 0.39 (i.e. the demonstrator freezes 39% of the time) if the observer does not freeze at all, plus 0.41 times the freezing of the observer if the observer does freeze. That the freezing of the observer was part of the model best explaining the data suggests that - unlike what a classic one-way perspective would assume - the behavior of the demonstrator is influenced by that of the observer. Indeed, the feedback parameters in the models all have 95% credibility intervals not encompassing zero, which provides additional evidence that the feedback was significant.

**Table 1.**
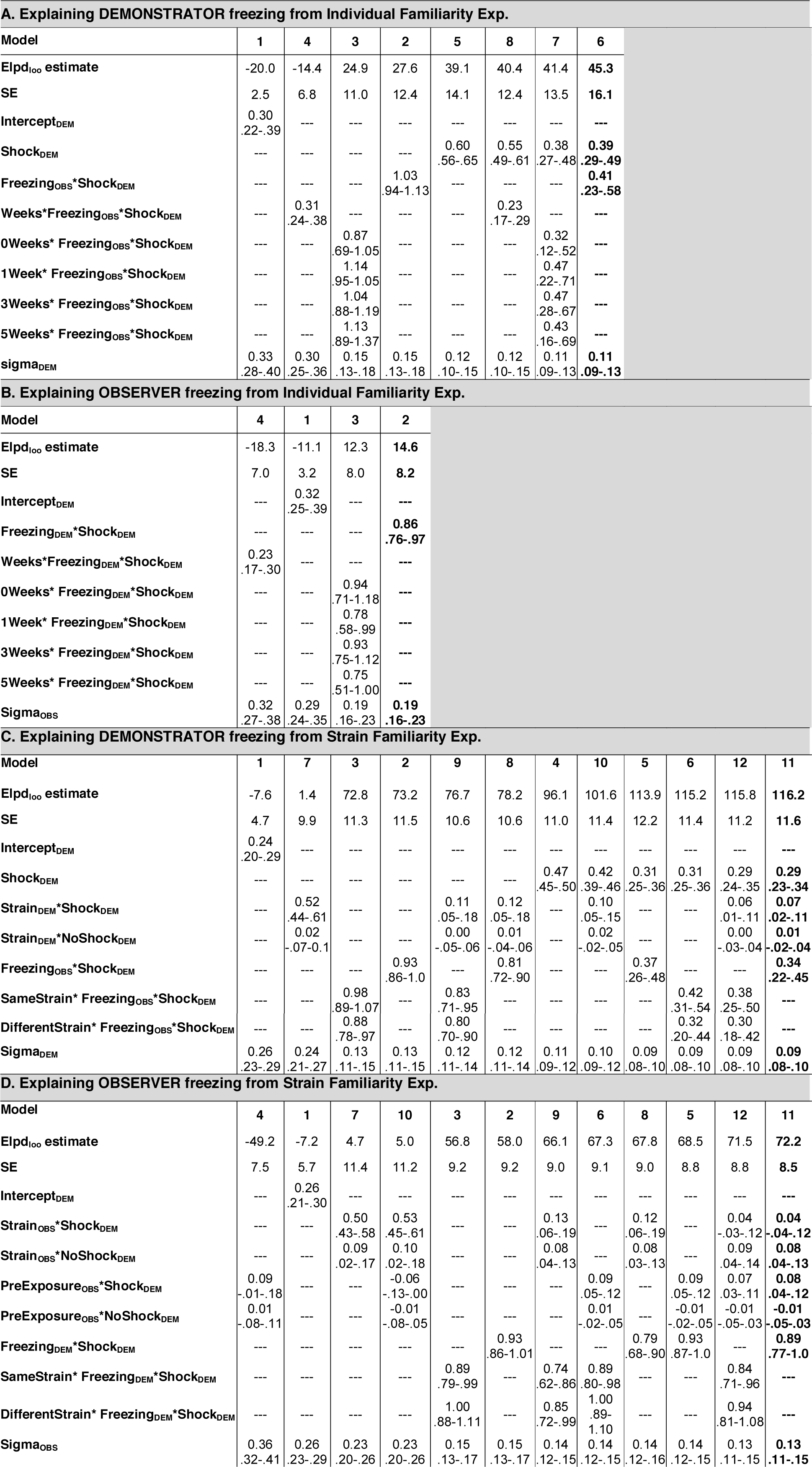
Model comparisons. For each experiment separately, separate models were constructed to describe the level of freezing of the demonstrator and the observer. The number of models varies depending on the variables that were included (Figure 4). The models were ordered based on their increasing leave-one-out predictive performance (ELPDloo) with the worst model left, and the best model right. The first column lists the variables included in each model. Values in the table indicate the parameter estimates with their credible interval below (from 2.5% - 97.5%). The last column in bold always indicates the winning model. Elpdloo: expected log pointwise predictive density according to the leave-one-out approximation. SE: standard error of the Elpdloo. DEM: demonstrator. OBS: observer.

As expected, delivery of footshocks is a key variable that induces freezing in the demonstrator. In contrast, none of the familiarity variables were present in the model with the best fit, indicating that familiarity does not modulate the freezing of demonstrators sufficiently to improve the predictive performance of the model. This was true independently of whether familiarity was modeled to vary linearly with weeks (model 8, where more weeks spent together would increase the influence of the other animal’s freezing) or non-linearly (model 7, where a different strength of influence from the observer freezing is calculated for each familiarity level). Indeed, inspecting the distribution of the parameters for the social feedback fitted separately for each group in model 7 (distributions in Figure 5A) shows substantial overlap between the credibility intervals for these parameters. Put differently, the data does not provide evidence that the freezing of an unfamiliar observer (0_weeks_) has a significantly smaller effect than that of more familiar observers. In addition, even for the 0_weeks_ group, the social feedback parameter has a distribution that is shifted away from zero, suggesting significant social feedback onto even unfamiliar demonstrators. Note, that in all models, the observer’s freezing was only considered as a predictor while the demonstrator received shocks (i.e. Freezing_obs_*Shock_dem_), and models that considered the freezing of the observer without the presence of a shock (e.g. during baseline) performed less well (data not shown).

**Figure 5.**
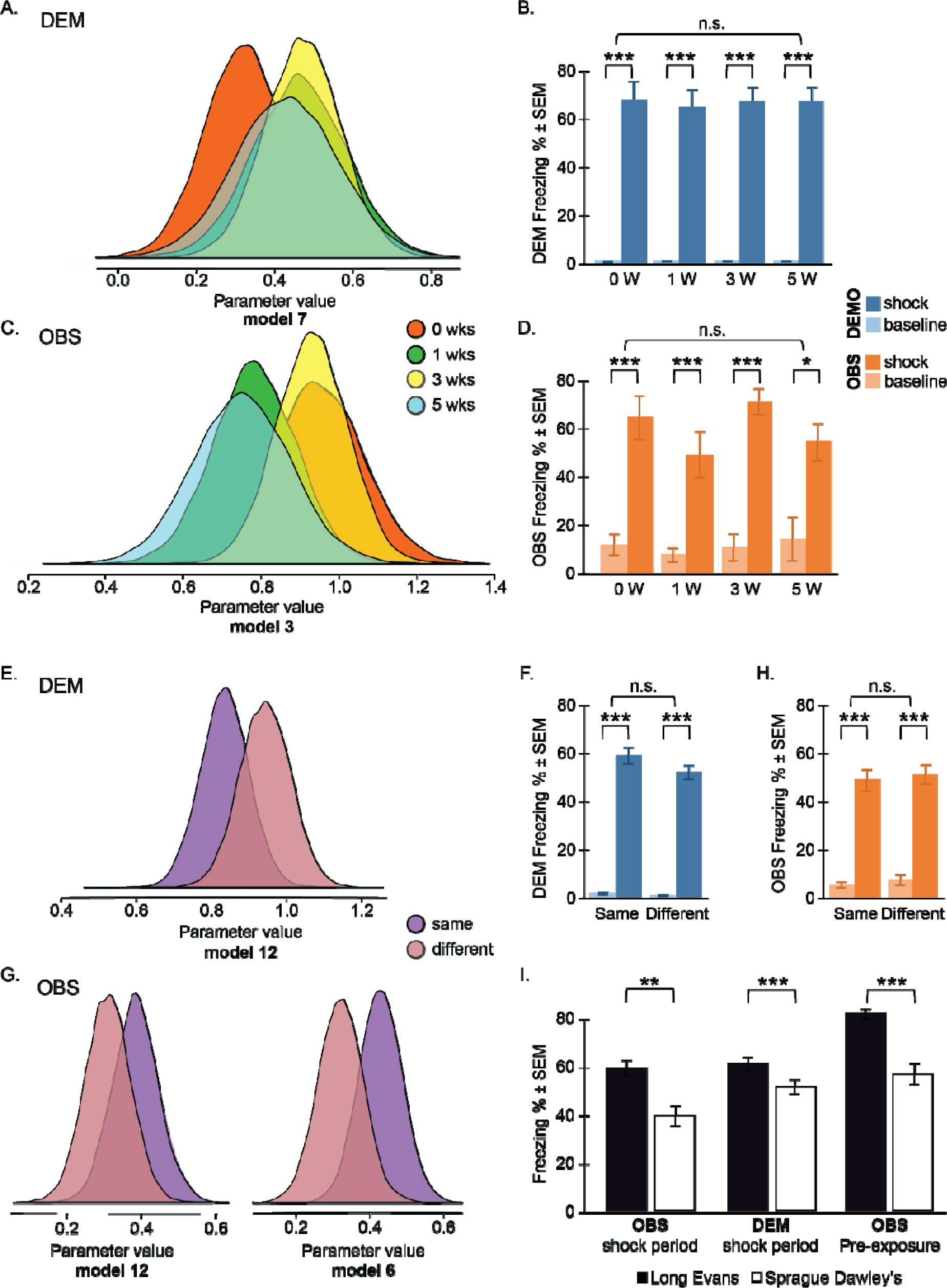
Parameter estimates and model-free analysis. (A) parameter estimates of the influence from OBS -> DEM from Model 7 in Table 1A as a function of weeks spent together. Note the considerable overlap and shift away from zero illustrating the lack of a familiarity effect and the consistent feed-back from the observer, respectively. (B) model free comparison across the familiarity groups. (C-D) same as A-B but using observer freezing as the dependent variable. E, F, G, H: same for the Strain Familiarity Experiment. I: Long Evans rats froze more than Spragues both in a social context during the shock period of the Strain Familiarity Experiment, and when tested alone during shock pre-exposure. For all pairwise comparisons, *t*-test, \**p*<0.05., \*\**p*<0.01, \*\*\**p*<0.001. NS: refers to the absence of a significant group x epoch interaction in an ANOVA (see text for details).

The findings from these model comparisons were confirmed with traditional group level analysis: the inclusion of Shock_dem_ as a significant parameter is reflected in a significant increase of freezing during shock compared to baseline for each group (Figure 5B) and the lack of familiarity effects is compatible with the outcome of a 2 epoch (baseline vs shock) x 4 familiarity (0, 1, 3, 5 weeks) ANOVA that revealed a main effect of epoch (F_(1,28)_=409.685, *p*<0.0001) but a lack of significant main effect of familiarity (F_(3,28)_=0.569, *p*=0.64) or familiarity x epoch interaction(F_(3,28)_=0.463, *p*=0.711).

##### Observer Freezing

Of the four defined models, the one best explaining the data estimated that Freezing_obs_ = 0.86 x Freezing_dem_ x Shock_dem_ (*model* 2, elpd_loo_ estimate = 14.6 and SE = 8.2, Table 1B). This shows that within a dyad the freezing of the observer (Freezing_obs_) is coupled to that of the demonstrator (Freezing_dem_) with a high gain of 0.86 x Freezing_dem_. In other words, the freezing of the observer is only 14% lower than that of the demonstrator receiving the actual shock. Inspecting the distribution of the parameters influenced by familiarity of model 3 reveals much overlap between the distributions, with all of them having credibility intervals not encompassing zero (Figure 5C). This further reinforces the notion that a strong linkage exists independently of the familiarity level. Traditional group level comparisons (Figure 5D) confirm that administering a shock to the demonstrator has a strong effect on the observer but that familiarity does not modulate this effect: a 2 epoch x 4 familiarity ANOVA showed a main effect of epoch (F_(1,28)_=113.069, *p*<0.0001), but no main effect of familiarity (F_(3,28)_=1.214, *p*=0.323) or epoch x familiarity interaction (F_(3,28)_=1.135, *p*=0.352).

Together the Bayesian Model Comparison on the Individual Familiarity experiment data therefore shows (a) that embracing a bidirectional model of emotional contagion improves our ability to explain the data and (b) that there is no apparent change in the intensity of the bi-directional coupling as a function of how long Long-Evans rats were pair-caged. The mutual influence evidenced here occurs for unfamiliar and familiar animals alike.

#### 2.2.2 Results from the Strain Familiarity Experiment

Unlike the Individual Familiarity experiment in which all animals were Long Evans, to further test the impact of familiarity, the Strain Familiarity experiment included rats of different strains: Long Evans and Sprague Dawley’s. The difference in strain could impact freezing in two ways. Much like in the first experiment, strain influenced how familiar the partners are with the strain of their counterpart: Long Evans rats were highly familiar with Long Evans rats but had never been in contact with Sprague Dawley rats. In addition, one of the two strains may in general freeze more to a stressor (be it social or non-social) than the other. This second consideration motivated us to include a number of factors in addition to those included in the first experiment (see Fig. 4). In addition to (1) the freezing percent of observers and demonstrators, and (2) whether or not the demonstrators received footshocks (baseline vs shock period), the following predictors were also included: (3) the strain of the observers as a predictor for observer freezing (Strain_obs_), (4) a binary variable capturing the strain (Long Evans or Sprague Dawley) of the demonstrators to predict demonstrator freezing (Strain_dem_), and (5) a binary variable that captured cases in which the two rats were of the same strain (SameStrain) and those in which they were of different strains (DifferentStrain). Finally, to capture individual differences in freezing behavior, which is crucial for predicting observer freezing, we also analyzed movies made during the initial pre-exposure of the observer rats in which they experienced a number of shocks alone, and used that as a predictor of how much they would respond to seeing another rat receive a shock (PE_OBS_) (Figure 3).

##### Demonstrator Freezing

The model best explaining the data estimated that: Freezing_dem_ = 0.29 x Shock_dem_ + 0.07 x Strain_dem_ x Shock_dem_ + 0.34 x Freezing_obs_ x Shock_dem_ (*model 11* elpd_loo_ estimate = 116.2 and SE = 11.6, Table 1C). This means that if no shock is being delivered, the estimated freezing is zero, because of the ‘x Shock_dem_’ behind all terms. If a shock is delivered, Freezing_dem_ is then estimated at 0.29 plus 0.07 if the demonstrator is a Long Evans plus 0.34 x the freezing of the observer. Examining the ranking of the models in Table 1C shows that as for the Individual Familiarity experiment, adding the presence of shock and feedback from the observer to predict the demonstrators freezing improved the fit to the data, but assuming that the effect of the demonstrator is different for same or different strains does not. The observers’ freezing was only a good predictor when the demonstrator actually received shocks (i.e. Frezing_obs_*Shock_dem_), and models that considered the freezing of the observer without the presence of a shock (e.g. during baseline) performed less well (data not shown).

Same and different strain variables overlap in their parameter distributions (Figure 5E), showing no difference in freezing of the demonstrator when paired with an observer of the same or different strain. The difference in freezing between strains during the shock period is confirmed by group level analyses showing Long Evans demonstrators froze significantly more compared to Sprague Dawley demonstrators (Figure 5I). Importantly, as for Experiment 1, the feedback parameters (i.e. those including Freezing_obs_) all had credibility intervals excluding zero, providing evidence for the presence of a sizable feedback effect. Additional model-free group level analyses (Figure 5F) confirm these findings: there was a significant increase of freezing levels during shock compared to baseline (epoch) and no effect of famil-iarity: a 2 epoch x 4 familiarity ANOVA showed a main effect of epoch (F_(1,58)_=637.323, p < 0.0001), but no main effect of familiarity (F_(1,58)_=2.491, p=0.12) or epoch x familiarity interaction (F_(1,58)_=1.695, p=0.198).

##### Observer Freezing

For the observer the model best explaining the data was: Freezing_obs_ = 0.08 x Strain_obs_ x Noshock_dem_ + 0.08 x Pre-exposure x Shock_dem_ + 0.89 x Freezing_dem_ x Shock_dem_ (*model 11*, elpd_loo_ estimate = 72.2 and SE = 8.5, Table 1D), showing that within a dyad the freezing of the observers (Freezing_obs_) during the shock period is strongly modulated by the freezing of the demonstrators (Freezing_dem_ x Shock_dem_) and more weakly by the pre-exposure of the observer (Pre-exposure_obs_*Shock_dem_). During the no shock period (i.e. baseline), the freezing of the observer is mildly modulated by the strain of the observer animal (Strain_obs_ x Noshock_dem_), which suggests possible differences between freezing levels of observers of different strains (Figure 5I). Whether observer-demonstrator dyads were from the same or different strains, however, did not modulate the strength of the coupling between demonstrator and observers’ freezing. An additional experiment which showed that Long Evan observers are capable of distinguishing same (other unfamiliar Long Evans) from different strain (unfamiliar Sprague Dawley rats) under red dim light conditions (i.e., same as in the Strain Familiarity experiment), confirmed that this lack of effect was not due to the possibility that the observers could not distinguish the two strains (Figure 6). This illustrates that the behavior of the observer is modulated by that of the demonstrator, regardless of whether they are from the same or different strain. This is further supported by an analysis showing no difference in freezing levels between same and different strain dyads (Figure 5H, left side): a 2 epoch x 4 familiarity ANOVA showed a main effect of epoch (F_(1,58)_=269.113, p < 0.0001), but no main effect of familiarity (F_(1,58)_=0.284, p=0.596) or epoch x familiarity interaction (F_(1,58)_=0.004, p=0.953).

**Figure 6.**
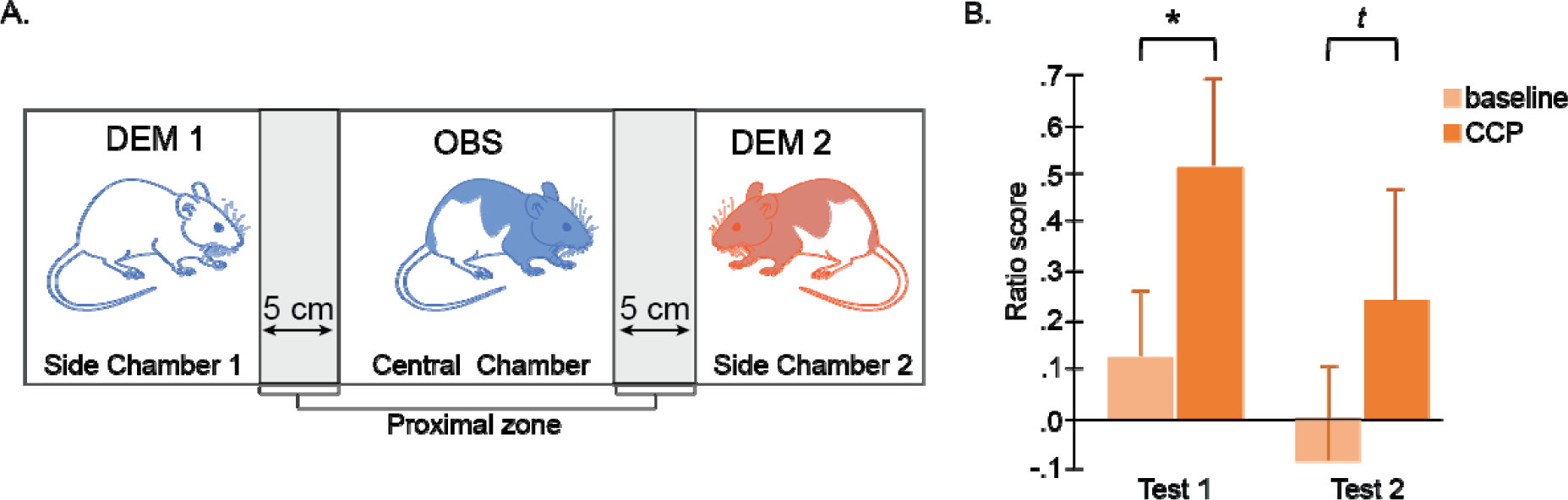
Same strain recognition experiment. For this experiment, eight observers (OBS: all Long Evans, four of which also served as demonstrators) and eight Demonstrators (DEM; four Long Evans and four Sprague Dawley rats) were used. A) The test was conducted in a three-chamber testing box consisting of one large central chamber (L:72cm x W:33cm) and two small side chambers (each: L:27cm x W:33cm). The central chamber was separated from the side chambers by transparent perforated walls. The day prior to test, all animals were habituated to the testing box. The test consisted of a 5 minute baseline period in which observers were individually placed in the central compartment, followed by a 10 minute choice preference period (CPP), in which two unfamiliar demonstrators (DEM1 and DEM2: one Long Evans and one Sprague Dawley rat) were simultaneously placed in one of the side compartments (placement was randomized). To avoid bias, the placement of the demonstrators occurred when the observer was in the center zone of the central compartment. Each observer had two tests in which the location of the Sprague Dawley and Long Evans rats was changed. The amount of time that the observers spent in the proximal zone during the initial 90 seconds of the baseline and CPP was scored and a ratio score was estimated (difference in the time spent in the proximal zone of the Long Evans rat and the time spent in the proximal zone of the Sprague Dawley rat divided by the sum of the time the observer spent in the proximal zone of the Long Evans and Sprague Dawley rat. B) Results show, that in test 1 and 2, observers spent more time in the proximal zone of the Long Evans demonstrators than that of the Sprague Dawley rats compared to baseline (paired sample t-test, one tail, *p =0.02 in test 1 and ^t^p=0.07 in test 2).

### Summary

Despite differences in experimental manipulations, both experiments suggest that there is robust bidirectional information transfer within observer-demonstrator dyads: (i) the freezing level of an observer is better predicted when taking the freezing of the demonstrator into account, (ii) the freezing level of a demonstrator is better predicted when taking the freezing of the observer into account and (iii) estimates of the coupling parameters have credibility intervals not including zero. In contrast, the familiarity level does not improve predictions, and the coupling parameters for different familiarity levels (individual or strain) overlap. This was true if familiarity was manipulated at the individual level in terms of weeks spent together or at the strain level in terms of whether animals were familiar with the strain of their partner.

### 2.3 Moment to moment emotional contagion - Granger causality

The results of Bayesian modeling provide evidence for bi-directional information transfer at the between dyad level. Dyads with higher overall observer freezing are dyads with higher demonstrator freezing despite receiving the same shock. If the freezing of the observer truly influences that of the demonstrator, as the Bayesian models suggest, we would expect to find evidence of such bi-directional influence at the level of the moment-to-moment fluctuations of freezing in individuals: fluctuations in the freezing of the demonstrator should be explained (in the statistical sense) by *earlier* fluctuations of the observer at a second by second time scale. Granger causality analyses were used to examine this prediction (Barnett & Seth, 2014; Seth, Barrett, & Barnett, 2015).

To have an overview of the information flow within the demonstrator-observer dyad during the emotional contagion test, Granger causality was computed including all dyads from both the Individual and Strain Familiarity experiments. G-causality values (i.e. Granger F values) were calculated separately for the baseline and the shock period. During baseline, significant G-causality was found in both directions: from demonstrator to observer (Granger *F* = 0.086, *p* < 0.0001) and from observer to demonstrator (Granger *F*= 0.156, *p* < 0.0001), meaning that there is time-coupled bidirectional information flow. The order of the Granger causality model is determined automatically by the analysis and was 21, suggesting that the freezing of an animal is influenced by the freezing levels in the past 21 seconds. In addition, the observer to demonstrator G-causality was numerically larger than in the opposite direction, which can be explained by the influence of the pre-exposure on the observer’s freezing: because the observers were pre-exposed to footshocks, they showed some contextual fear generalization to the test setup and more spontaneous freezing during the baseline (Figure 3, black marginal histograms). This baseline freezing potentially influenced the demonstrators. Conversely, as the demonstrators froze less during baseline, they could not have as much influence on the observers. For the shock period, significant G-causality was also found in both directions: from demonstrator to observer (Granger *F*= 0.059, *p* < 0.0001) and from observer to demonstrator (Granger *F*= 0.035, *p* < 0.0001). As expected, delivery of foot shocks to the demonstrator makes the information flow from the demonstrator to the observer stronger compared to the opposite direction, as indicated by the larger G-causality value in the demonstrator to observer direction.

To investigate the effect of familiarity, G-causality between the demonstrator’s and the observer’s freezing was calculated for each dyad in each direction (i.e. demonstrator to observer and *vice versa*) separately, and then compared between different experimental groups. Due to the fact that during baseline both demonstrators and observers showed minimal freezing levels, there were not enough freezing time points to calculate the G-causality for each dyad during this period. Therefore, to examine the effect of familiarity the analysis was restricted to the shock period (Figure 7A).

**Figure 7.**
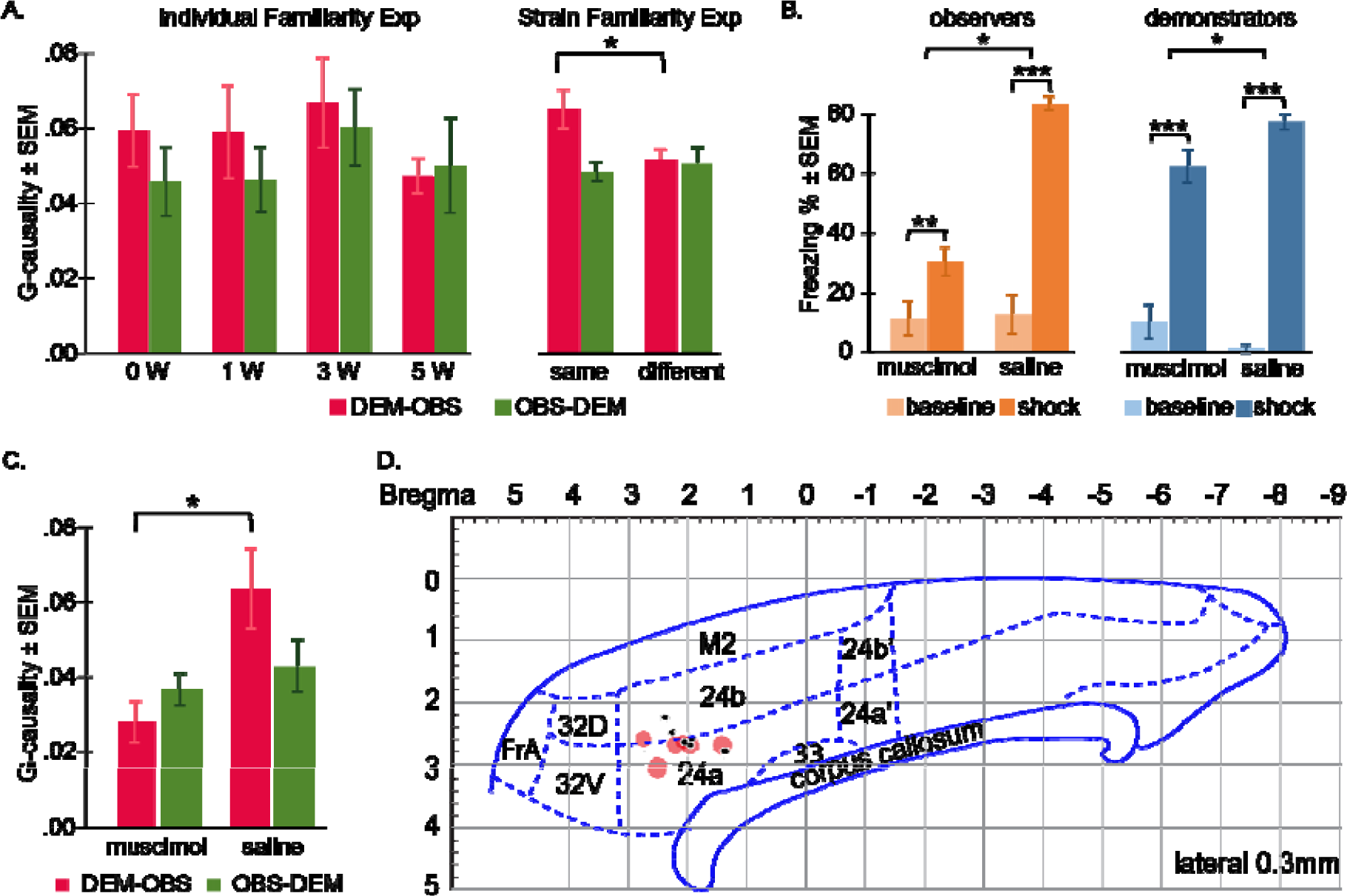
Directionality of emotional contagion and the role of the ACC. (A) G-causality results. Mean ± standard error of the mean (SEM) for G-causality values for the demonstrator to observer direction (DEM-OBS, in red) and the observer to demonstrator direction (OBS-DEM, in green) during the shock period, for both the Individual (left) and Strain (right) Familiarity experiments. W: week, same=same strain, different=different strain. (B) Effect of ACC deactivation on freezing. Percentage of time the observers (in orange) and demonstrators (in blue) spent freezing during baseline (light color) and the shock period (dark color) after ACC deactivation (muscimol) or after control treatment (saline). Freezing % = 100*freezing time/total time of the corresponding period. (C) Effect of ACC deactivation on the flow of information. Mean ± standard error of the mean (SEM) of the G-causality values in the demonstrator to observer direction (DEM-OBS, in red) and in the observer to demonstrator direction (OBS-DEM, in green) during the shock period, after ACC deactivation (muscimol) or after control treatment (saline). (D) Localization of the deactivations. Location of saline (black dots) and muscimol injections (red circles) on a sagittal view of the rat cortex [adapted from The Rat Brain in Stereotaxic Coordinates, 7th Edition, Paxinos and Watson]. The surface of each red circle is proportional to the z-score of the freezing level of that animal relative to the average and standard deviation of the control group (m=83% of time freezing, s=6.8%). Coordinates for each animal were determined by estimating the location of the tip of the cannula from coronal Nissl stainings, and averaging the estimate of the right and left cannula.

#### Individual familiarity

A MANOVA with G-causality of both directions as dependent variables and familiarity (0, 1, 3 & 5 weeks) as fixed factors revealed no significant effect of familiarity in either direction: (1) demonstrator to observer (*F* _*(3,28)*_ = 0.437, *p* = 0.728) and (2) observer to demonstrator (*F*_*(3,28)*_ = 0.496, *p* = 0.688), indicating that time spent together as cagemates did not affect the temporal coupling of the freezing of the dyad (Figure 7A).

#### Strain familiarity

A MANOVA with G-causality of both directions as dependent variables and familiarity (same strain versus different strains) as fixed factors revealed a small effect of condition in the demonstrator to observer direction (*F*_*(1,58)*_ = 4.726, *p* = 0.034) but not in the observer to demonstrator direction (*F*_*(1,58)*_ = 0.210, *p* = 0.648). In the demonstrator to observer direction, the G-causality was bigger for same-strain dyads compared to dyads composed of different strains, indicating that there was more information flow from the demonstrator to the observer when both animals were from the same strain than when they were from different strains (Figure 7A)

### 2.4 Role of the Anterior cingulate cortex (ACC) in emotional contagion

Given that both model comparison and granger causality suggest that the behavior of the observer feeds back on the behavior of the demonstrator, we wanted to experimentally probe this feedback by reducing the freezing reaction of the observer and testing whether that would reduce freezing in the demonstrator. In humans, the ACC has been considered one of the core regions activated by witnessing the pain of others (Bernhardt & Singer, 2012; Keysers, Kaas, & Gazzola, 2010; Lamm, Decety, & Singer, 2011). This region has its homologue in the ACC of the rat (Vogt, 2014) and has been implicated in emotional contagion and empathy in rodents as well (Allsop et al., 2018; Burkett et al., 2016; de Waal & Preston, 2017; Jeon et al., 2010; Keysers & Gazzola, 2017; B. S. Kim et al., 2014; S. Kim et al., 2012). We therefore predicted that deactivating this region in observers should reduce their vicarious freezing and, by virtue of the feedback connection the Individual and Strain Familiarity experiments suggest, reduce the freezing of the demonstrator. To examine this possibility and confirm the role of the ACC in social information transfer in rodents, a third experiment was conducted in which the ACC of the observers was deactivated using muscimol, and the impact on vicarious freezing was studied in both observers and demonstrators (note that this condition is part of a larger experiment available here https://doi.org/10.1101/450643).

#### Effect of muscimol on observer freezing

A 2 (periods: baseline vs shock) x 2 (condition: muscimol vs saline groups) repeated measures ANOVA was conducted to test the effect of ACC deactivation on socially triggered freezing of the observers (Figure 7B). All observers froze significantly more during the shock period (mean ± SD=30.24% ±11.77% for the muscimol and mean ± SD=83.57% ± 6.79% for the saline group) than during the baseline (mean ± SD=11.21% ± 14.35% for the muscimol and mean ± SD=12.60% ± 18.87% for the saline group) as confirmed by the significant main effect of period (baseline vs shock: *F*_*(1,12)*_=126.556, *p*<0.0001;). Paired sample t-tests confirmed that in both conditions the observers’ freezing levels were significantly higher during the shock period compared to the baseline (muscimol group: *t*_*(5)*_=5.617, *p*<0.005; control group: *t*_*(7)*_ =11.101, *p*<0.0001) showing socially triggered freezing in both ACC-deactivated and control observers. However, observers with ACC-muscimol injection froze significantly less compared to saline controls (main effect of condition: *F*_*(1,13)*_=100.805, *p*< 0.0001) indicating that the ACC is necessary for full-fledged socially triggered freezing. A significant period x condition interaction effect was also found (*F*_*(1,13)*_=31.737, *p*<0.0001) reflecting that the impact of muscimol was larger during the shock period.

#### Effect of muscimol on demonstrator freezing

To test our hypothesis that demonstrators paired with muscimol observers would show reduced freezing compared to those paired with saline observers, a one tailed t-test was performed on demonstrator freezing during the shock period and results were significant (*t*_*(12)*_=2.397, *p*<0.024; Figure 7B). An ANOVA including condition (muscimol vs saline) x epoch (baseline vs shock) confirmed this effect as a significant interaction (*F*_*(1,12)*_=19.837, *p*< 0.001), with the effect of condition larger during the shock than baseline.

#### Granger-Causality

To further investigate the impact of ACC deactivation on the temporal coupling across the animals, a Granger analysis was performed on the second-to-second freezing of the observers and the demonstrators (Figure 7C). It was expected that deactivating the ACC of the observer should perturb the information transfer from the demonstrator to the observer, because a structure necessary for triggering vicarious freezing in the observer (i.e. the ACC) would be impaired. It was also expected that the transfer in the observer to demonstrator direction should remain unaffected because the brain of the demonstrator was not injected with muscimol. To compare differences between the two groups, a MANOVA with G-causality of each dyad in both directions (demonstrator-observer and observer-demonstrator) as dependent variables and conditions (muscimol versus saline) as fixed factors was conducted. A significant effect of condition in the demonstrator to observer direction (*F*_*(1,12)*_=6.620, *p*=0.024) was found but not in the observer to demonstrator direction (*F*_*(1,12)*_=0.424, *p*=0.527). In the demonstrator to observer direction, the G-causality was significantly smaller for the ACC-deactivated group compared to control dyads, indicating that the observers’ freezing responses were less influenced by the demonstrators’ when the observers’ ACC were deactivated, and that the temporal dynamic within the dyad was impaired by the manipulation.

#### Histological Reconstruction

histological reconstructions confirmed that we successfully targeted the ACC, particularly region 24a and 24b (Figure 7D).

### 2.5 Danger detection interpretation – Computational modeling approach

A surprising outcome of the results was that emotional contagion seems to be mutual even in unfamiliar animals. Why would an animal exposed to electroshocks modulate its expressions of distress based on the reaction of unfamiliar bystanders, and why should a bystander care about the pain of strangers? If the primary purpose of emotional contagion were to generate empathy and promote prosocial behavior, one would expect that it would increase with higher familiarity and affiliation (de Waal & Preston, 2017). Here we therefore explored the utility of emotional contagion towards a more selfish motive: danger assessment. We designed simulations that explore whether in the presence of uncertainty, including the behavioral reaction of others, can improve the accuracy of danger detection.

Several simulations were performed that compared danger detection performance of individuals with or without social information (i.e. taking or not taking the freezing from another animal into account), and with equal or unequal access to the danger signals (see methods for details). Briefly, the logic of the simulations is that a danger signal is turned on and off over time (blue in Figure 8A), generating an internal danger signal in the animal after addition of noise of magnitude σ. In the individual condition, the animal then decides whether to freeze or not to freeze based on whether the internal signal surpasses a threshold (yellow in Figure 8A), leading to a time series of freezing decisions (red in Figure 8A and time series shown in Figure 8B). In the social simulation, the individual additionally takes into account the freezing at time t-1 of the other animal in deciding whether to freeze at t, by adding b*(freezing_other(t-1)_- 0.5) to its internal danger signal (Figure 8B).

**Figure 8.**
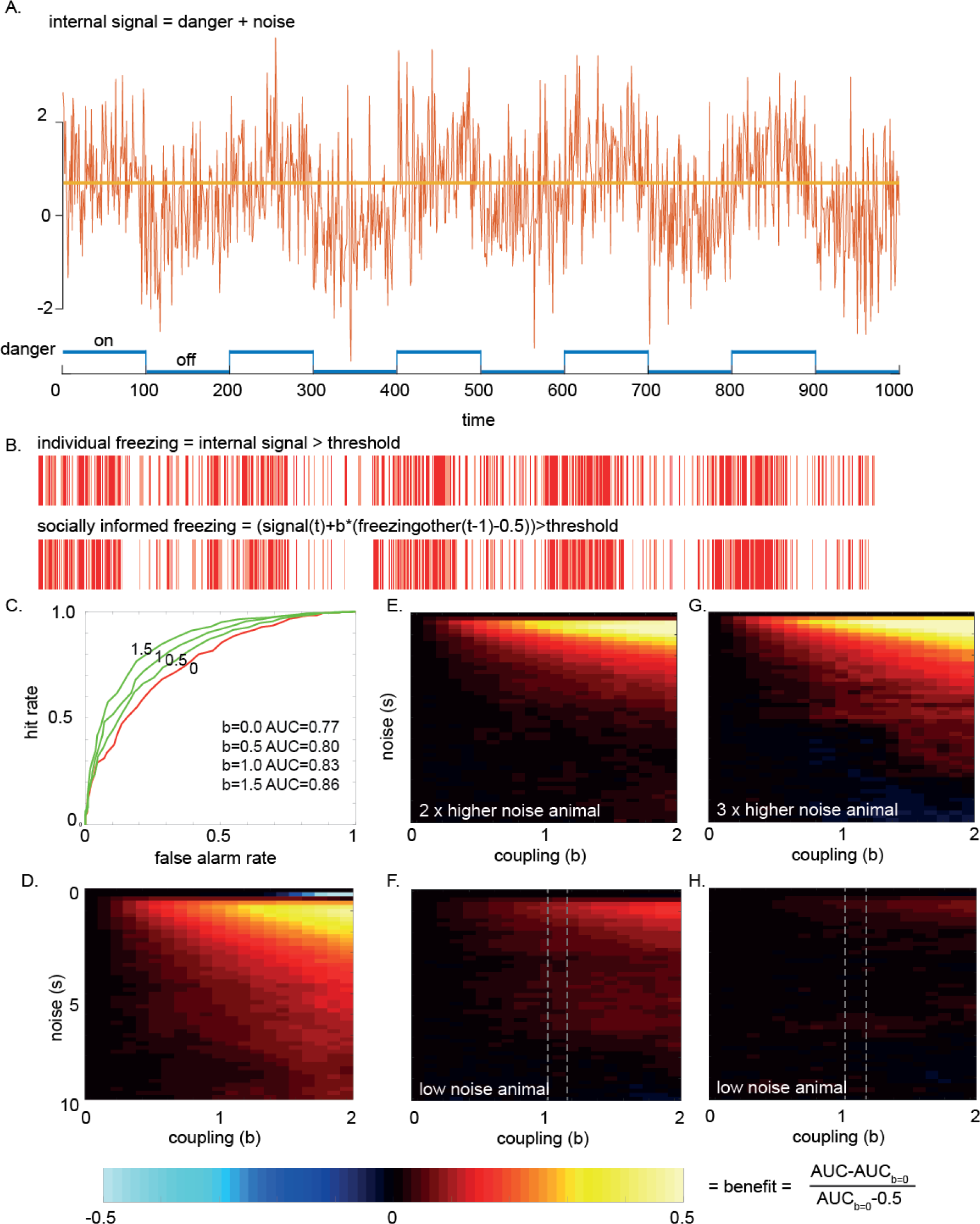
Computational modeling supports a danger detection interpretation. (A) Internal danger signal simulated for an animal that is exposed to 100 timepoints of danger and 100 timepoints of no danger with noise added. The animal freezes when the danger signal surpassed a certain threshold (yellow line). (B) Time series of freezing for an animal by itself (individual freezing) or for one that is additionally taking the freezing of another animal into account (socially informed freezing). (C) Accuracy of danger detection shown as the area under the ROC curve (AUC) for different coupling factors (b=0 to 1.5). A higher coupling factor increases the AUC. (D) Benefit of taking the freezing of others into account when both animals have the same access to danger signals, i.e. experience the same noise level. (E-H) Animals with twice (E) or thrice (G) as much noise as compared to another animal (F and H, respectively) had stronger benefits from coupling. However, the low noise animals (F and H) experience no disadvantages. The dotted lines indicate the coupling regime that our animals appeared to be in the Individual and Strain Familiarity experiments.

When both animals have the same access to danger signals (i.e. experience the same signal to noise ratio), the decision to freeze becomes more accurate if animals take the freezing of the other animal into account. Figure 8C illustrates this phenomenon at relatively low noise level (σ=1). If the animal does not take the freezing of the other into account (coupling b=0, red curve), the area under the red receiving operating characteristic curve (AUC) is equal to 0.77. Increasingly taking the freezing of the other into account (b from 0.5 to 1.5) augments the AUC, meaning a more accurate danger detection. This benefit in danger detection can be seen as comparatively more freezing (Figure 8B, red) when danger is present and less when it is absent for the socially informed freezing. This means that animals that are influenced by the freezing of the other will freeze more when there is danger and freeze less when there is none. Repeating this analysis for different noise levels (σ) and coupling (b) reveals that over a wide range of parameters, there are either benefits (red and yellow colors) or no disadvantage (black) (Figure 8D). Only in very specific cases (very low noise σ<0.5 and high coupling b>1) is there a loss of performance (Figure 8D). Analyzing the time series (data not shown) shows that these rare cases occur when the animal that is no longer in danger (timet) erroneously persists in its freezing because the other animal was freezing at t-1.

In our experiments, one animal however has privileged access to danger signals because it experiences the shock itself, while the other has less direct access. What was surprising is that the more informed demonstrators still relied on the behavior of the less informed observers. To examine such scenarios additional models were simulated to capture unequal access to danger signals. This was done by imposing twice or thrice as much noise on one animal compared to the other. In these models the animal with more noise has stronger benefits from coupling (Figure 8E and G) however, the other animal experiences no disadvantages (no cold color in Figure 8H) and sometimes even advantaged (warm colors in Figure 8F).

One may wonder how these coupling parameters compare to those we found in our Bayesian models for the demonstrators. In our simulation, b represents the ratio of the social/direct danger signal. Accordingly, in our Bayesian models for demonstrator freezing (Table 1), it can be approximated as the fraction freezing_obs_/shock_dem_, and would have the value b=1.05 and b=1.17 (gray dotted lines in Figure 8F and H) for the Individual and strain Familiarity experiment, respectively.

In summary, we find that moderate coupling in the order of magnitude found in our Bayesian modeling (b∼1) always improved the decision-making of our simulated animals.

## 3. Discussion

Various studies in recent years have provided evidence for social information transfer within rodent dyads (for review see Keum et al., 2016; Meyza et al., 2017; Panksepp & Lahvis, 2011; Sivaselvachandran et al., 2016). Even though different kinds of paradigms were developed, they were all restricted to testing the impact of one stimulus animal on a target animal. For instance, in observational freezing paradigms (Atsak et al., 2011; Bredy & Barad, 2009; Gonzalez-Liencres et al., 2014; Jeon et al., 2010; Keum et al., 2016; Kiyokawa, Honda, et al., 2014; Knapska et al., 2010; Langford et al., 2006) the stimulus animal receives foot shocks or is exposed to a pre-conditioned cue while the target animal passively witnesses this, allowing researchers to examine the impact from the stimulus animal to the target animal (Figure 1A). In contrast, in social buffering paradigms the target animal is subjected to distress while being witnessed by a stimulus animal that can be in different (stressful or non-stressful) states, allowing the other direction to be measured (Fuzzo et al., 2015; Ishii et al., 2016; Kikusui et al., 2006; Kiyokawa, Kikusui, Takeuchi, & Mori, 2004; Mikami et al., 2016) (Figure 1B). However, the measurements in these paradigms were always restricted to one direction and the social impact in the other direction seldom received attention (for an exception, see Langford et al., 2006). This unidirectional approach potentially results in an impoverished understanding of social interactions.

The core aims of our experiments were twofold. First, to explore whether influences in social transmission paradigms should be recognized as bidirectional. Second, whether familiarity indeed has the effect on these influences that is often assumed in the literature. To do so, we leveraged established analysis techniques from other fields to analyse our rodent distress transmission paradigm. These methods include Bayesian model comparison, which is prominent in for example neuroeconomics (e.g. Glimcher & Fehr, 2013) and, in addition, Granger causality analyses, which are used in neuroscience to examine information transfer across neural populations (Seth et al., 2015).

Bayesian modeling revealed that there was not only information transfer from the demonstrator to the observer rat but also feedback from the observer to the demonstrator rat. This was even the case across unfamiliar individual and unfamiliar strains. Granger causality analyses further confirmed temporal coupling between the demonstrator and the observer in both directions. To our knowledge, this is the first rigorous quantitative demonstration of bi-directional social information transfer in the now widely used rodent emotional contagion paradigms and provides a better fit to the data than the traditional one-way focus of current studies. Conceiving of the influence as mutual has the conceptual advantage of integrating social distress transmission and social buffering as two sides of the same mechanism, thereby providing a unifying framework across related fields that have so far engaged in relatively little cross-talk.

In terms of neural mechanisms, we show that the ACC is crucially involved in this mechanism. More precisely, temporarily deactivating this brain structure in one member of the social interaction attenuates the information transfer to the injected individual. Furthermore, this deficit feeds back and influences the behavior of the brain-intact partner, showing again bi-directional information transfer.

This finding raises the question of why dyads of rats should engage in bidirectional emotional contagion. Is there an evolutionary benefit? Traditionally, our interpretation of social distress transmission has been shaped by the belief that this phenomenon depends on familiarity. Whereas there is substantial evidence that familiarity influences this phenomenon in mice, this has not systematically been investigated in experiments with rats. Our results show that although it may be intuitive to extrapolate the effectiveness of a factor from mice to rats, this is not the case for familiarity. Critically, across our experiments, we find bidirectional information transfer across unfamiliar animals. This is true across Long Evans rats that have never been housed together and across different strains of rats that have never witnessed members of the other strain. Accordingly, familiarity is certainly not a pre-requisite for emotional contagion in rats. The difference in social structure between mice and rats may account for part of this difference (Lund, 1975). Mice do not tolerate other mature males around them, and the presence of other mature individuals thus triggers a strong stress reaction per se, that inhibits information transfer (Martin et al., 2015). Rats, on the other hand, live in much larger groups with other adult males that are tolerated (Lund, 1975). It may be that in that structure, seeing an unfamiliar individual does not produce the kind of stress response that would shut down information transfer, and thereby allows the significant emotional contagion we document in our design. An alternative explanation is that the rats showed distress contagion in all cases because they failed to recognize the difference between familiar and unfamiliar partners. However this cannot account for our data since our control experiment (Fig. 6) demonstrates that the rats can perceive the difference between familiar and unfamiliar individuals at the illumination levels used during our emotional contagion paradigm.

Not only do we find that information transfer is significant in unfamiliar animals and strains, we also find comparatively little evidence that the information transfer is increased in more familiar animals. The Bayesian model fitting shows that the parameter estimates for the freezing transmission is similar across the different familiarity levels, with largely overlapping distributions. The Bayesian model comparison further shows that models that stratify the connection based on familiarity do not outperform models that assume the same strength of connections for all groups. These models, however, were calculated based on the overall level of freezing in the entire 12 minute period. It is possible that the effect of familiarity is more evident in a fine-grained analysis of the second to second decision to freeze. However, in the Individual Familiarity experiment, such a fine-grained Granger causality analysis also evidenced no effect of familiarity, while confirming a significant coupling across all dyads. That is to say, dyads that saw each other for the first time on the day of testing coordinated their freezing as closely as those that had spent five weeks together. Only towards extreme strangers, i.e. animals of a strain they had never encountered before, did that analysis reveal a small decrease of granger causality, and then only in the demonstrator towards observer direction. In other words, although observers will respond to the shock given to the demonstrator of an unkown strain (with levels of freezing similar to those when witnessing their own strain, as revealed by the Bayesian modeling), the moment at which they will show that reaction is slightly less tightly linked to that of the demonstrator compared to animals from the same strain.

The bidirectional information flow we demonstrate and the weak effects of familiarity in rats we observe are difficult to reconcile with the notion that emotional contagion, as measured using vicarious freezing, is *primarily* a mechanisms for empathy, that directs prosocial behavior to kin (de Waal & Preston, 2017; Preston & de Waal, 2002). Instead, our computer simulations argue for a simpler interpretation that sees this mechanism as a means to compute danger signals in a crowd. We demonstrate, using simplified simulations, that the accuracy of danger detection in a noisy environment is improved if an animal takes the freezing behavior of other animals into account. Importantly, in the parameter range that we find in our Bayesian modeling, in which demonstrators give similar weights to the shock and social information, we found that taking social information into account never decreases the danger detection performance of the simulated individuals in a group. Whereas these simulations have many limitations, in particular the fact that they assume that noise is independent across animals, they encourage us to think of emotional contagion and social buffering not so much as mechanisms meant to benefit others, but as mechanisms that social animals should develop in order to improve their own danger detection. Picking up the emotions of others becomes akin to using others as antennas to amplify often noisy danger signals. In this interpretation, emotional contagion (Figure 1A) then occurs when the behavior of the another animal signals higher danger whereas social buffering (Figure 1B) happens when the behavior of another animals indicates lower danger. At a group level, the dyad comes to a consensus on the level of danger, something that has been shown to improve decision-making and has motivated the field of crowd decision-making (Dyer et al., 2008). This per-spective does not imply that emotional contagion cannot serve empathy and prosocial decision making. Rather, it suggests that the strong familiarity gating observed for prosocial motivations in rats (Ben-Ami Bartal et al., 2014) is likely to occur after emotional contagion has happened, at least in rats. Emotional contagion itself may be more neutral, and serve the crowd computation of danger, which needs not to be gated by familiarity. Traditionally, eavesdropping, in particular the fact that some mammals and birds show signs of fear when they perceive the alarm-calls of *other* species (Magrath, Haff, Fallow, & Radford, 2015) has been conceived of as different from emotional contagion across individuals of the *same* species. The former (which has to our knowledge received little attention in rats) is considered a selfish form of information gathering while the latter has been seen as more prosocial. Our data invites us to consider that they may not be as different after all, and invite us to investigate in the future, the role structures such as the ACC have in eavesdropping.

## Materials and methods

### Subjects

For experiment 1 (i.e., Individual familiarity), 128 male Long Evans rats (6-8 weeks old), for Experiment 2 (i.e., Strain familiarity), 164 male rats (83 Long Evans and 81 Sprague Dawleys /6-8 weeks old) and for experiment 3 (i.e., deactivation of the ACC, also reported in https://doi.org/10.1101/450643), 60 male Long Evans rats were all obtained from Janvier Labs (France). Upon arrival animals were housed in groups of 4 or 5. Only animals of the same strain were housed in the same cage. All animals were maintained at ambient room temperature (22-24oC, 55% relative humidity, SPF, type III cages, on a reversed 12:12 light-dark cycle: lights off at 07:00) and allowed to acclimate to the colony room for 7 days. Food and water were provided *ad libitum*. All experimental procedures were pre-approved by the Centrale Commissie Dierproeven of the Netherlands (AVD801002015105) and/or by the welfare body of the Netherlands institute for Neuroscience (IVD, protocol number NIN151101, NIN1493 and NIN151104).

### Setup

All tests were conducted in a two-chamber apparatus (each chamber L: 24cm x W: 25m x H: 34cm, Med Associates, Inc.). Each chamber consisted of transparent Plexiglas walls and stainless steel grid rods. The compartments were divided by a transparent perforated Plexiglas separation, which allowed animals in both chambers to see, smell, touch and hear each other. For shock pre-exposure of observers and for the emotional contagion tests one of the chambers was electrically connected to a stimulus scrambler (ENV-414S, Med Associates Inc.). For video recording of the rats’ behaviors, a Basler GigE camera (acA1300-60gm) was mounted on top of the apparatus controlled by EthoVision XT (Noldus, the Netherlands).

### Experimental procedures

All experimental procedures for Experiments 1 and 2 were conducted during the first 5 hours of the dark part of the day light cycle. Figure 2 illustrates the general procedures used for both experiments.

### Experiment 1 –Individual famliarity

#### Experimental groups

Observer-demonstrator dyads were randomly allocated to one of the following groups: unfamiliar condition (n=7 dyads), familiar for 1 week (n=10 dyads), familiar for 3 weeks (n=9 dyads) or familiar for 5 weeks (n= 7 dyads).

#### Handling and habituation

Prior to the start of the experimental procedures, animals were randomly paired and assigned the role of observer or demonstrator. Depending on the familiarity condition, observer-demonstrator dyads were housed for 1, 3 or 5 weeks prior to test. For the unfamiliar condition, 3 weeks prior to test day, animals were housed in dyads of the same role (i.e. either two observers or two demonstrators in one cage). Ten days prior to the emotional contagion test, all animals were handled every other day for 3 minutes. To habituate animals to the testing conditions, four days preceding testing, animal dyads were transported and placed in the testing apparatus for 20 min/day for three consecutive daily sessions. The testing apparatus was cleaned with lemon-scented dishwashing soap and 70% alcohol in between each dyad.

#### Shock pre-exposure

To enhance the emotional contagion response to the distress of the demonstrators (Atsak et al., 2011), observer animals experienced a shock pre-exposure session the day prior to test day. The shock pre-exposure was conducted in one of the chambers of the test apparatus. To prevent contextual fear, the walls of the chamber were coated with black and white striped paper, the background music was turned off, the apparatus was illuminated with bright white light and the chamber was cleaned with rose-scented dishwashing soap and vanilla aroma drops. Observers were individually placed in the apparatus and after a 10 minute baseline, four footshocks (each: 0.8mA, 1 sec long, 240-360sec random inter-shock interval) were delivered. After the shock pre-exposure session, animals were placed for 1 hour in a neutral cage prior to return to their home cage.

#### Emotional contagion test

The testing setup was illuminated with dim red light,cleaned using a lemon-scented dish-washing soap followed by 70% alcohol, and background radio music was turned on. Each observer-demonstrator dyad was transported to the testing room and animals were placed in the corresponding chamber of the testing apparatus. For the unfamiliar condition, randomly chosen observers and demonstrators from different cages were used to create the testing dyads. For this condition, observers and demonstrators never had contact with each other until test start. For the familiar conditions, observers and demonstrators were from the same cage. The testing order was fully randomized. For all dyads, following a 12 minute baseline, the demonstrators experienced five footshocks (each: 1.5mA, 1 sec long, 120 or 180sec inter-shock interval). Following the last shock, dyads were left in the apparatus for 2 additional minutes prior to returning to their home cage.

### Experiment 2-Strain familiarity

#### Experimental groups

Observer-demonstrator dyads were randomly allocated to one of four groups in which the demonstrators received footshocks in the emotional contagion test and two control groups in which no shocks were delivered during the test. The experimental groups consisted of; 1) dyads of two Long Evans (LE-LE; n=19 dyads), 2) dyads of two Sprague Dawleys (SD-SD; n=13 dyads), 3) dyads of a Long Evans observer and a Sprague Dawley demonstrator (LE-SD; n= 17 dyads) or 4) dyads of a Sprague Dawley observer and a Long Evans demonstrator (SD-LE; n= 11 dyads). The control groups included dyads of two Sprague Dawleys (SD-SD-no-shock; n=5 dyads) and dyads of a Sprague Dawley observer and a Long Evans demonstrator (SD-LE-no-shock; n= 17 dyads).

#### Handling and habituation

Upon arrival, all animals were randomly paired in same-strain and same-role dyads (i.e., each dyad of animals was assigned the role of either observer or demonstrator), which were pair-housed together. Handling and habituation procedures were conducted in the same way as in experiment 1 with the exception that the shock pre-exposure was conducted following the first habituation and this was followed by the second and third habituation sessions. In addition, during habituation a white plastic perforated floor was added on top of the grid floor of the observer’s chamber.

#### Shock pre-exposure

The shock pre-exposure for all animals was conducted following the first habituation session. The shock pre-exposure parameters were identical to those described for experiment 1.

#### Emotional contagion test

The testing procedures and parameters for experiment 2 were the same as those described for the unfamiliar condition of experiment 1. Observers and demonstrators were randomly chosen according to the experimental condition (e.g. for the SD-LE condition a Sprague Dawley from an observer cage and a Long Evans from a demonstrator cage were selected). Although all animals were kept in the same room during acclimation, observers and demonstrators did not have contact with each other (nor to any individual of a different strain) until the start of the test. Similar to habituation, a white perforated plastic was placed on top of the grid floor of the observer’s chamber.

### Experiment 3-ACC deactivation

#### Note

The shock observation condition reported here is part of a larger experiment reported in https://doi.org/10.1101/450643.

#### Experimental groups

Observer-demonstrator pairs were randomly allocated to one of two groups: saline control group (n=10) or muscimol group (n =8). Four dyads (2 from control group and 2 from muscimol group) were excluded after histology examination suggesting damage of corpus callosum due to injection.

#### Handling

Upon arrival, all animals were randomly housed in dyads, one assigned as the observer and one as the demonstrator.

#### Surgery and guide-cannula implantation

Cannulas were implanted into the ACC 1 week prior to behavioral testing (hit: n=14; miss: n=4). All animals were anesthetized with isoflurane (1-3%). The animals were then positioned in a stereotaxic frame with blunt-tipped ear bars, and a midline incision was made. Six burr holes were drilled (2 for anchoring screws and 1 for the cannula per hemisphere). Two single guide-cannulas (62001; RWD Life Science Co., Ltd) were implanted targeting bilateral ACC (AP, +1.7; ML, ±1.6; DV, +3.5 mm with a 20° angle from the surface of the skull, Paxinos and Watson, 1998) and chronically attached in the observer animals with a thin layer of acrylic cement (Super-Bond C & B ®, Sun Medical Co. Ltd., Shiga, Japan) and thick layers of acrylic cement (Simplex Rapid, Kemdent, UK). To prevent clogging of the guide cannula, a dummy cannula (62101; RWD Life Science Co., Ltd) was inserted and secured until the microinjection was administered.

#### Habituation

Habituation procedures were conducted in the same way as for experiment 1 and 2 except that prior to transport to the experimental room, the observer animals were habituated to a sham infusion procedure.

#### Microinjections

Fifteen minutes prior to the emotional contagion test, observer animals were lightly restrained, the stylet was removed and an injection cannula (62201; RWD Life Science Co., Ltd) extending 0.8 mm below the guide cannula was inserted. Muscimol (0.1 <u>μ</u>g/μl) or saline (0.9%) was microinjected using a 10 μl syringe (Hamilton), which was attached to the injection cannula by PE 20 tubing (BTPE-20; Instech Laboratories, Inc.). A volume of 0.5 μl per side was injected using a syringe pump (70-3007D; Harvard Apparatus Co.) over a 60 s period, and the injection cannula remained untouched for an additional 60 s to allow for proper absorption and to minimize pull up effect along the track of the cannula. The protective cap was secured to the observer animal after the infusion and then the animal was returned to its home cage.

#### Shock pre-exposure

The shock pre-exposure for all animals was conducted following the first habituation session. The shock pre-exposure parameters were identical to those described for experiment 1. All shocks during the shock pre-exposure were co-terminated with a tone stimulus (2.5 kHz, around 70db, 20seconds). This tone was then played back to the animals on a later day in a control experiment that is not further reported here.

#### Emotional contagion test

The testing procedures and parameters for experiment 3 were the same as those described for the familiar condition of experiment 1. Similar to habituation, a white perforated plastic was placed on top of the grid floor of the observer’s chamber.

### Behavior scoring

The behavior of observers and demonstrators during the emotional contagion test and/or pre exposure was manually scored by 2 experienced researchers (inter-rater reliability assessed with Pearson’s r correlation coefficient was > 0.9) and using the open source Behavioral Observation Research Interactive Software (BORIS, Friard & Gamba, 2016). Freezing, defined as lack of movement except for breathing, was continuously scored throughout the 12 minutes baseline and 12 minutes shock period. To create a continuous time series, freezing moments extracted from the Boris result files were recoded as 1 and non-freezing moments as 0 using Matlab (MathWorks inc., USA) on a second to second basis. For experiment 3 (i.e., deactivation of the ACC), the researcher that scored the movies was blind to the experimental manipulation (i.e., control or muscimol group).

### Statistics

#### General linear models

The results of experiments 1, 2 and 3 and the freezing responses of observers and demonstrators were analyzed separately. Freezing time was calculated as the sum of all freezing moments in a certain epoch and freezing percentage was calculated as the total freezing time divided by the total time of the epoch. Baseline period (1st epoch) was defined as the first 710-seconds of the emotional contagion test and the shock period (2nd epoch) was defined as the 710-second following the first shock (approx. 720 seconds from the start of the test). For comparison between periods and conditions, repeated measures ANOVAs (IBMSPSS statistics, USA) were performed with baseline and shock period as within subject factors and the conditions were used as between-subject factors (Experiment 1: 0,1,3,5 weeks; Experiment 2: same strain dyads, different strain dyads; Experiment 3: saline group, muscimol group.

### Bayesian Model Estimation and Comparison

For experiment 1, models were designed using combinations of the following variables: the freezing percent of observers and demonstrators, the number of weeks that demonstrator-observer dyads were housed together (0, 1, 3 and 5 weeks) and whether or not the demonstrators received footshocks (baseline vs shock period). For experiment 2, models were designed using all possible different combinations of the following variables: the freezing percent of observers and demonstrators, whether demonstrator-observer pairs were from the same (Long Evans – Long Evans, Sprague Dawley – Sprague Dawley) or different strain (Long Evans – Sprague Dawley, Sprague Dawley – Long Evans), whether or not the demonstrators received footshocks (baseline vs shock period), the freezing percent of the observers during pre-exposure and the strain of the observers and demonstrators (Long Evans or Sprague Dawley).

Note, that in all cases, we only considered the freezing of the other animal during the shock period, by multiplying them with the dummy variable Shock_dem_ that had a value of zero during baseline and one when a shock was applied. This was done for two reasons. First, our previous experiments had shown that prior shock experience was necessary for emotional contagion to occur in our paradigm (Atsak et al., 2011), and for the demonstrators, this prior experience was only available after the first shock. Second, inspection of the data (Figure 2) confirmed that the relation between observer and demonstrator freezing that is apparent during the shock period (red) was not apparent during the baseline period (black) where there seemed to be a disconnect between large individual variance in observer freezing (y-axis) and much smaller variance in demonstrator freezing (x-axis). We used relatively flat priors for all parameters with a normal distribution of mean 0 and standard deviation 2. The parameters were initially restricted to real numbers ranging from −1 to 1. For the link between observer and demonstrator freezing we noticed that estimates sometimes got close to 1. For those parameters we then relaxed the range to −1.5 to 1.5, and results in the table stem from these less constrained bounds.

Model fitting and parameter estimation were conducted using Bayesian analysis by estimating the posterior distribution through Bayes rule using in-house code in R Stan (Development Team, 2016) in R version 3.3.2 (R Core Team, 2016). All models converged (Rhat =1). To evaluate the predictive accuracy of each model a leave-one-out cross-validation (PSIS-LOO) was used to estimate the pointwise-out-of-sample prediction accuracy (elpd_loo) from all the fitted Bayesian models using the log-likelihood evaluated at the posterior simulation of the parameter values(A Vehtari, Gelman, & Gabry, 2016; Aki Vehtari, Gelman, & Gabry, 2016). To select a winning model, models were ranked based on their elpd_loo estimate and the model with the highest fit was compared pairwise to each of the other models until a first significant difference from the best model was reached. Specifically, we used the function ‘compare’ from the ‘loo’ library, and considered a difference significant, if the difference was at least one standard error away from zero. Amongst the winning models (i.e. those not significantly different from the one with the highest elpd_loo), the one with the highest fit was chosen as the winning model.

### Granger

Granger causality is a statistical concept of causality that is based on prediction (Granger, 1969). If a signal X1 “granger-causes” (or “g-causes”) a signal X2, then past values of X1 should contain information that helps predict X2 above and beyond the information contained in past values of X2 alone. In this study, X1 and X2 were binary time series of freezing of the demonstrator and freezing of the observer (freezing coded as 1 and not-freezing coded as 0) on a second-to-second basis. The freezing of the observer at a certain time point (*X2(t)*) can be estimated either by its own history plus a prediction error (reduced model, 1) or also including the history of the freezing of the demonstrator (full model, 2).

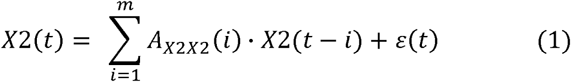

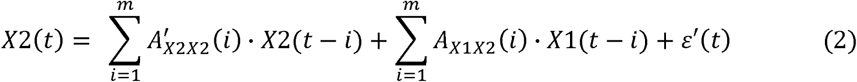

In equations 1 and 2, *t* indicates the different time points (in steps of 1s), *A* represents the regression coefficients and *m* refers to the model order which is the length of the history included. Granger causality from the freezing of the demonstrator to the freezing of the observer (i.e. X1→X2) is estimated by comparing the full model (2) to the reduced model (1). Mathematically, the log likelihood of the two models (i.e. G-causality value *F*) is calculated as the natural logarithm of the ratio of the residual covariance matrices of the two models (3).

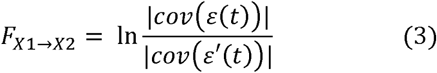

This G-causality magnitude has a natural interpretation in terms of information-theoretic bits-per-unit-time (Barnett & Seth, 2014). In this study, for example, when G-causality from the demonstrator to the observer reaches significance, it indicates that the demonstrator’s freezing can predict the observer’s freezing and that there is information flow from the demonstrator to the observer. Jumping responses of the demonstrator to the foot shocks were also taken into account and a binary time series of this behavior was included as X3 (jumping coded as 1 and not-jumping coded as 0). Given that the demonstrators did not exhibit any jumping during baseline, X3 was only included in the analysis done on the shock period.

The algorithms of the Multivariate Granger Causality (MVGC) Toolbox (Barnett & Seth, 2014) in MATLAB were used to estimate the magnitude of the G-causality values. First, the freezing time series of the demonstrators and the observers were smoothed with a Gaussian filter (size = 300s, sigma =1.5). The MVGC toolbox confirmed that each time series passed the stationary assumption for Granger causality analysis. Then, the optimal model order (*m*, the length of history included) was determined by the Akaike information criterion (AIC) for the model including all observer-demonstrator dyads. The optimal model order is a balance between maximizing goodness of fit and minimizing the number of coefficients (length of the time series) being estimated. For experiment 1 and 2, the model order of 21 was estimated to be the best fit for the model including all dyads and thus it was fixed at 21 for the subsequent dyad-wise analysis. The largest model order across all dyads was 22 and running the analysis by fixing the model order to 22 showed similar results. For experiment 3, the estimated best model order was 19 and thus it was fixed at 19 for the dyad-wise analysis. To test the differences of the G-causality values across conditions, multivariate ANOVAs were performed using SPSS.

### Simulations

The logic behind the simulations was to explore the hit and false alarm rate of two individuals in a dyad that take decisions to freeze or not to freeze based on an internal danger signal that results from an objective danger signal plus noise. A given time-point was considered a hit if the animal froze and danger was present, and a false alarm if the animal froze but the danger was absent. Two cases were compared: one where there is no information exchange between animals (individual case), and one where there is information flow between animals (social case). In both cases, an animals’ internal danger signal was triggered by witnessing a danger signal d(*t*) that was on for 100 time-samples then off for 100 time samples for 5 cycles for a total of 1000 time points (see equation 1)

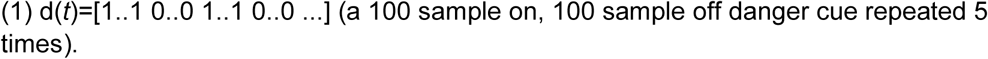

Both animals experienced noise on top of the signal, with the noise being independent across animals (equation 2). Noise level was varied systematically by changing σ:

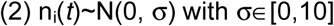

In the no feedback model, the internal signal of each animal *i* was simply the addition of signal and noise (equation 3):

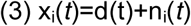

And animals decide to freeze or not to freeze based on whether the signal is above or below threshold c (equation 4):

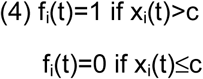

In the model with feedback and equal access to the danger signal, we aimed to simulate a situation in which both animals have a similar access to the danger signal but are also sensitive to the freezing of the other animals. We thus calculated the internal signal iteratively by additionally considering whether the other animal froze on the preceding time-point or not, with both animals experiencing equal noise levels. The degree to which the internal signal depends on the freezing of the other is systematically varied using b∈[0,2]. Given that both the danger signal and the freezing of the other animal take on values of zero and 1, b=1 means that the animal pays equal attention to sensory and social sources of information. (equation 5).

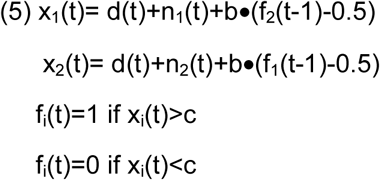

Finally, in models with feedback but unequal access to the danger signal, we aimed to simulate conditions in which one animal has more access to the danger signal than the other by adding r times more noise to animal 1 than 2. In that case, the degree to which the two animals consider the freezing from the other is scaled based on experienced noise, with animals experiencing more noise paying more attention to the freezing in the other (equation 6). This decision was informed by our finding that demonstrators are less influenced by observers than vice versa, and by the finding that humans integrate the influence of others in similar ways (Bahrami, Olsen, Latham, Roepstorff, & Frith, 2012).

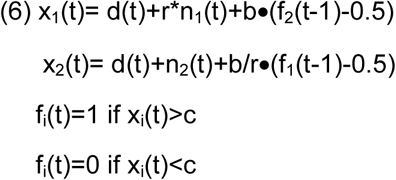

Performance was measured based on signal detection theory as the area under the ROC curve. Specifically, c is varied systematically from −5σ to +5σ, and the hit and false alarm rate is calculated in each case, with a hit being a freezing decision when the danger signal was 1, and a false alarm when it was 0. These rates are then plotted on an ROC curve, with false alarm as x and hit as y coordinates. Random decisions lead to AUC (area under the curve) of 0.5, perfect decisions to AUC=1. The gain in performance between the individual and social condition was calculated as (AUCsocial-AUCindividual)/(AUCindividual-0.5) to express how much further from chance the performance has become.

To explore more systematically the influence of noise level (σ), coupling (b) and noise ratio (r), for each combination of parameters we calculated performance gains 20 times (using new random numbers for the noise), and display the median of these 20 random noise sets.

## Acknowledgements

This work was supported by the Netherlands Organization for Scientific Research (VICI: 453-15-009 to C.K. and VIDI 452-14-015 to VG) and the European Research Council of the European Commission (ERC-StG-312511 to C.K.). We thank Michael Spezio for helpful discussions on the project, in particular on model comparison and for suggesting to use rstan and LOO approaches.

## Author Contributions

CK and VG conceived the study, supervised the team and acquired the funding. YH, RB and MC refine the experimental design. YH, RB, VP, NJ, MH, IB, SV, IP, TvL acquired the behavioral data; VP, NJ, MH, IB, SV, IP, TvL, NC rated the behavior with training from YH, RB and MC. MC performed the day-to-day supervision of the data acquisition. RT and YH performed the Granger analysis with guidance from CK. RB and CK performed the Bayesian model comparisons. CK performed and analysed the simulations. YH, RB, VG and CK wrote the first draft of the paper while all authors participated in revising the draft.

## Competing interests

Authors declare no competing interests.

## Data and materials availability

Data will be made available upon reasonable request.

